# Kismet/CHD7/CHD8 affects gut biomechanics, the gut microbiome, and gut-brain axis in *Drosophila melanogaster*

**DOI:** 10.1101/2021.12.17.473216

**Authors:** Angelo Niosi, Nguyên Henry Võ, Punithavathi Sundar, Chloe Welch, Aliyah Penn, Yelena Yuldasheva, Adam Alfareh, Kaitlin Rausch, Takhmina Rukhsar, Jeffery Cavanaugh, Prince Yadav, Stephanie Peterson, Raina Brown, Alain Hu, Any Ardon-Castro, Darren Nguyen, Robert Crawford, Wendy Lee, Mikkel Herholdt Jensen, Eliza J. Morris, Kimberly Mulligan

## Abstract

The gut-brain axis may contribute to the pathophysiology of neurodevelopmental disorders, yet it is often unclear how risk genes associated with these disorders affect gut physiology in a manner that could impact microbial colonization. We addressed this question using *Drosophila melanogaster* with a null mutation in *kismet,* the ortholog of chromodomain helicase DNA-binding protein (*CHD*) family members *CHD7* and *CHD8.* In humans, *CHD7* and *CHD8* are risk genes for neurodevelopmental disorders with co-occurring gastrointestinal symptoms. We found *kismet* mutant flies have a significant increase in gastrointestinal transit time, indicating functional homology of *kismet* with *CHD7/CHD8* in vertebrates. To measure gut tissue mechanics, we used a high-precision force transducer and length controller, capable of measuring forces to micro-Newton precision, which revealed significant changes in the mechanics of *kismet* mutant guts, in terms of elasticity, strain stiffening, and tensile strength.
Using 16S rRNA metagenomic sequencing, we also found *kismet* mutants have reduced diversity of gut microbiota at every taxonomic level and an increase in pathogenic taxa. To investigate the connection between the gut microbiome and behavior, we depleted gut microbiota in *kismet* mutant and control flies and measured courtship behavior. Depletion of gut microbiota rescued courtship defects of *kismet* mutant flies, indicating a connection between gut microbiota and behavior. In striking contrast, depletion of gut microbiome in the control strain reduced courtship activity. This result demonstrated that antibiotic treatment can have differential impacts on behavior that may depend on the status of microbial dysbiosis in the gut prior to depletion. We propose that Kismet influences multiple gastrointestinal phenotypes that contribute to the gut-brain axis to influence behavior. Based on our results, we also suggest that gut tissue mechanics should be considered as an element in the gut-brain communication loop, both influenced by and potentially influencing the gut microbiome and neuronal development.

## Introduction

The symbiotic relationships we share with our microbiome are critical for human development and adult homeostasis (1). The gut-brain axis specifically refers to the communication loop that exists between the gut microbiome and brain. Manipulation of gut microbiota can impact neurodevelopment and neurological function (2–4). In the opposite direction, brain-targeted interventions like cognitive behavioral therapy can modulate the gut microbiome (5). Studies seeking to define the molecular mediators of microbiota-gut-brain crosstalk have identified a variety of key players, including serotonin (6), short-chain fatty acids (SFCAs) (7), and lipopolysaccharides (8), which can communicate through the vagus nerve system (9–12).

Mounting evidence indicates that the gut microbiome is an etiological factor of neurodevelopmental disorders (NDDs) (13, 14). Analysis of fecal content has demonstrated that people with autism spectrum disorder (ASD) have altered gut microbiota when compared to neurotypical controls (14–17). Among individuals with ASD, common features of microbial dysbiosis in the gut include reduced microbial diversity and altered abundance of the predominant phyla, Firmicutes and Bacteroidetes (14, 16–20). Treating the gut dysbiosis of children with ASD with fecal microbiota transplant (FMT) from neurotypical donors can improve symptomatic behaviors (15). This same phenomenon is observed in mice; FMT from a wild-type mouse to a mouse model of ASD improved behavioral outcomes in the recipient, whereas FMT from the ASD mouse model to a wild-type mouse induced autism-like behaviors (21). Further, administration of FMT in mice using stool samples from humans with ASD caused behavioral impairments in the recipient mice, suggesting that similar types of microbial dysbiosis can elicit behavioral deficits across host species (22).

Determining how genes associated with NDDs affect gut physiology and microbial colonization could expand treatment options for both gastrointestinal (GI) discomfort and behavioral symptoms. *Drosophila melanogaster* are increasingly being used to examine the gut-brain axis given the relative simplicity of their tissues and gut microbiome (23), combined with the conservation of intestinal pathophysiology between flies and mammals (24). Fruit flies also possess orthologs to risk genes associated with NDDs, including *kismet,* the ortholog to mammalian chromodomain helicase DNA-binding domain protein (CHD) family members, *CHD7* and *CHD8.* In humans, mutations in *CHD7* cause a congenital NDD called CHARGE syndrome (25), and *CHD8* is among the highest confidence risk genes for ASD (26–28). Both CHARGE syndrome and *CHD8*-associated ASD have co-occurring GI abnormalities, including reduced gut motility and constipation (29, 30).

In *Drosophila, kismet* is broadly expressed in the developing brain (31), as well as in intestinal stem cells (32) and enteroendocrine cells (33). Neurodevelopmental and behavioral phenotypes attributed to Kismet include axon growth and guidance (31), axon pruning (31, 34), synaptic vesicle recycling (35), synaptic transmission (36), sleep (37), locomotion (20), and memory recall (34, 37). Kismet is also critical for maintaining intestinal stem cell homeostasis (32), though its role in the gut has not been fully elucidated.

Here, we show that *Drosophila* with heterozygous loss of *kismet* exhibit a range of GI phenotypes. The *kismet* mutants had a slower GI transit time and distinct gut tissue mechanics, including changes in elasticity, strain stiffening, and tensile strength. Analysis of the gut microbiome revealed that *kismet* mutants had an altered abundance of multiple bacterial taxa in both the anterior and posterior midguts, including a decrease in Firmicutes and an increase in opportunistic pathogens. Depletion of gut microbiota using streptomycin increased courtship activity of *kismet* mutant flies, indicating a connection between the *kismet* mutant*-*associated gut microbiota and behavior. In contrast, depletion of gut microbiota in the control strain induced courtship defects, demonstrating that microbial depletion can have variable impacts on behavior that likely depend on the level of gut dysbiosis. We propose that *kismet* partially influences the gut-brain axis by affecting interconnected aspects of gut physiology—GI transit time, biomechanics, and microbial composition—though further investigation is needed to delineate the reciprocal interplay and molecular underpinnings of the observed phenotypes. Additionally, we suggest that mechanical communication pathways are a critical component of the gut-brain axis.

## Materials and Methods

### Fly husbandry

Flies were reared on a standard cornmeal-yeast-agar medium recipe that was adapted from a Bloomington *Drosophila* Stock Center recipe. All flies were maintained at 23°C, except flies used for courtship analysis, which were maintained at 25°C in a humidified incubator on a 12:12 hour light-dark cycle. The *kismet (kis)* LM27 mutant strain —a generous gift from Dr. Daniel R. Marenda (Drexel University, Philadelphia, PA)—has a null allele of *kis* created by ethyl methanesulfonate (EMS) mutagenesis (38). Because homozygous null *kis/kis* is embryonic lethal, we used heterozygous *kis^LM27^* mutant flies with a CyO balancer to maintain the null allele. To create an isogenic control strain, *kis/CyO* were outcrossed to a balancer strain (+/CyO) in which the + chromosome was marked by *Scutoid.* The two strains were intercrossed for ten generations and the resulting *kis*/CyO and +/CyO were used for all analyses. Canton S flies, used for courtship analysis, were from the Bloomington Stock Center.

### Gastrointestinal transit time

Male flies aged 1-7 days post-eclosion were starved for 24 hours in hydrated starvation vials to ensure empty bowels and to induce hunger. After 24 hours, flies were placed in individual observation tubes (clear straws cut into thirds) containing food colored blue with 0.5% Bromophenol Blue, based on the method used by (39). Once blue food was ingested, indicated by blue food in the abdomen, we began recording time. Flies were repeatedly observed in five-minute increments until blue excrement was observed.

### Midgut length

Gastrointestinal tracts of male flies aged 1-7 days post-eclosion were removed in PBS using a dissecting microscope. Digital images were captured using a Motic dissecting microscope outfitted with a digital camera. ImageJ was used to measure midgut length.

### Biomechanical measurements

The full-length gut was dissected from flies aged 1-7 days post-eclosion and immediately mounted between two clips (Aurora Scientific, Aurora, ON) in PBS, which were attached to either side of the midgut. The clips were mounted to suspend the gut between a 322C-I High-Speed Length Controller and a 403B Force Transducer (Aurora Scientific, Aurora, ON). The initially slack gut was pulled along its length at a rate of 0.01 mm/s until breaking while monitoring the tissue’s extension, Δ*L*, and tensile force, *F*. During pulling, the samples were imaged with the 10X objective lens of a standard dissection microscope. All force and extension data were collected using LabVIEW (National Instruments, Austin, TX) and analyzed in Matlab (Mathworks, Natick, MA). The extension of the gut was normalized by its initial length, *L*, as the strain: 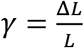. The linear stiffness of the tissue was determined as the slope of the force-strain curve at 0% strain (i.e., for an unstretched gut), while the maximal stiffness was determined as the maximal slope of the force-strain curve during the pull (Figure 2A). The tensile strength was quantified as the maximal force and strain the tissue could reach before breaking.

**Figure 1.**
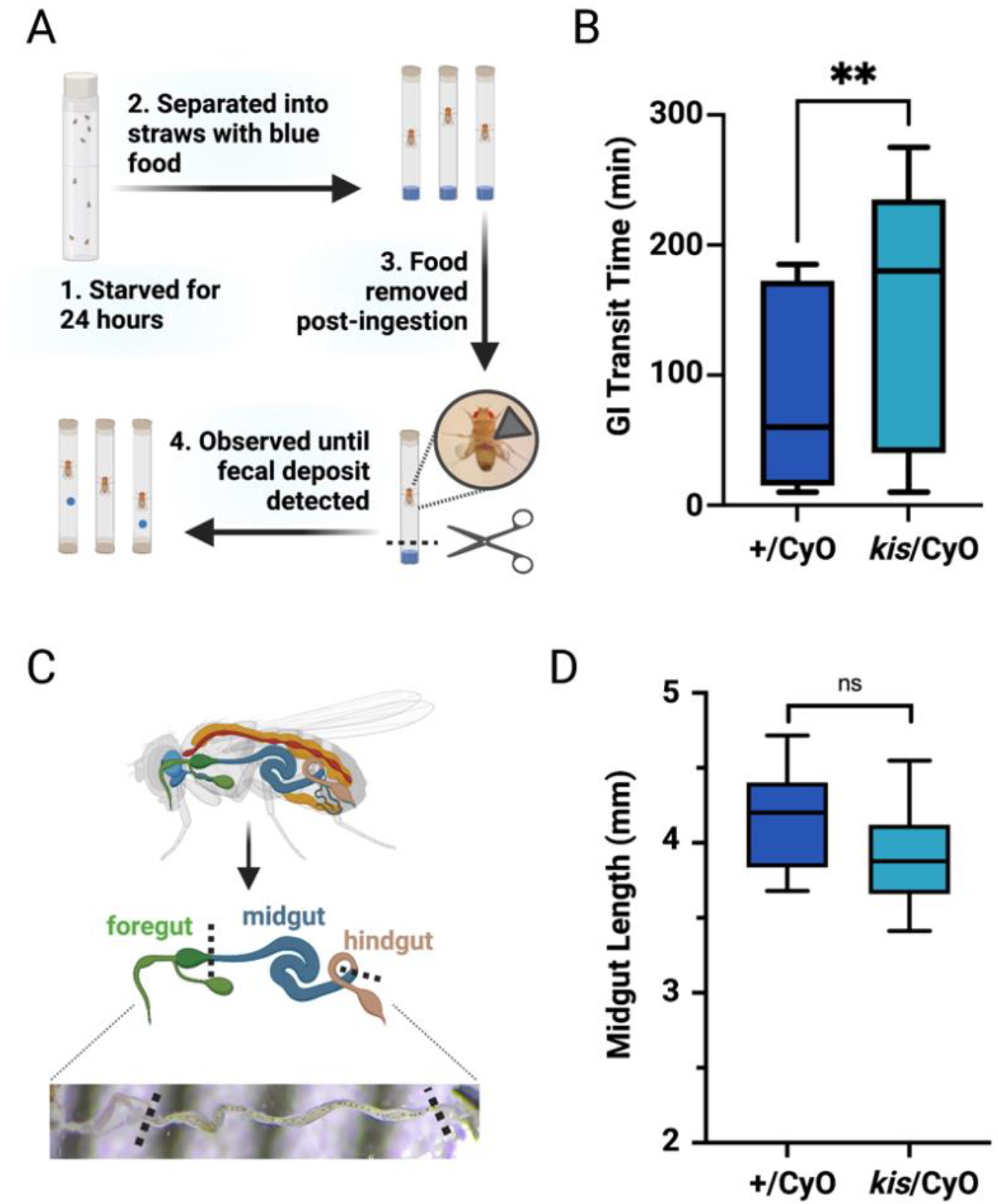
Kismet affects GI transit time but not midgut length. (A) Experimental scheme for measuring GI transit rate. Following a 24-hour starvation, flies were separated into individual straws containing blue food. Following ingestion, which was detected by the presence of blue food in the abdomen, the food was removed. Flies were monitored constantly until a blue fecal deposit was detected. (B) *kis*/CyO flies had a significantly slower GI transit time compared to control (+/CyO) flies. Mann-Whitney U test; ** p < 0.01; n = 17 for +/CyO, n = 23 for *kis/CyO*. (C) Gut dissection and measurement scheme. Dissected midguts were measured using ImageJ. Dashed lines indicate measurement boundaries. (D) Midgut lengths were not significantly different between control and *kis/CyO* flies. Student’s *t*-test; ns = not significant; n = 10 for +/CyO, n = 10 for *kis*/CyO.

**Figure 2.**
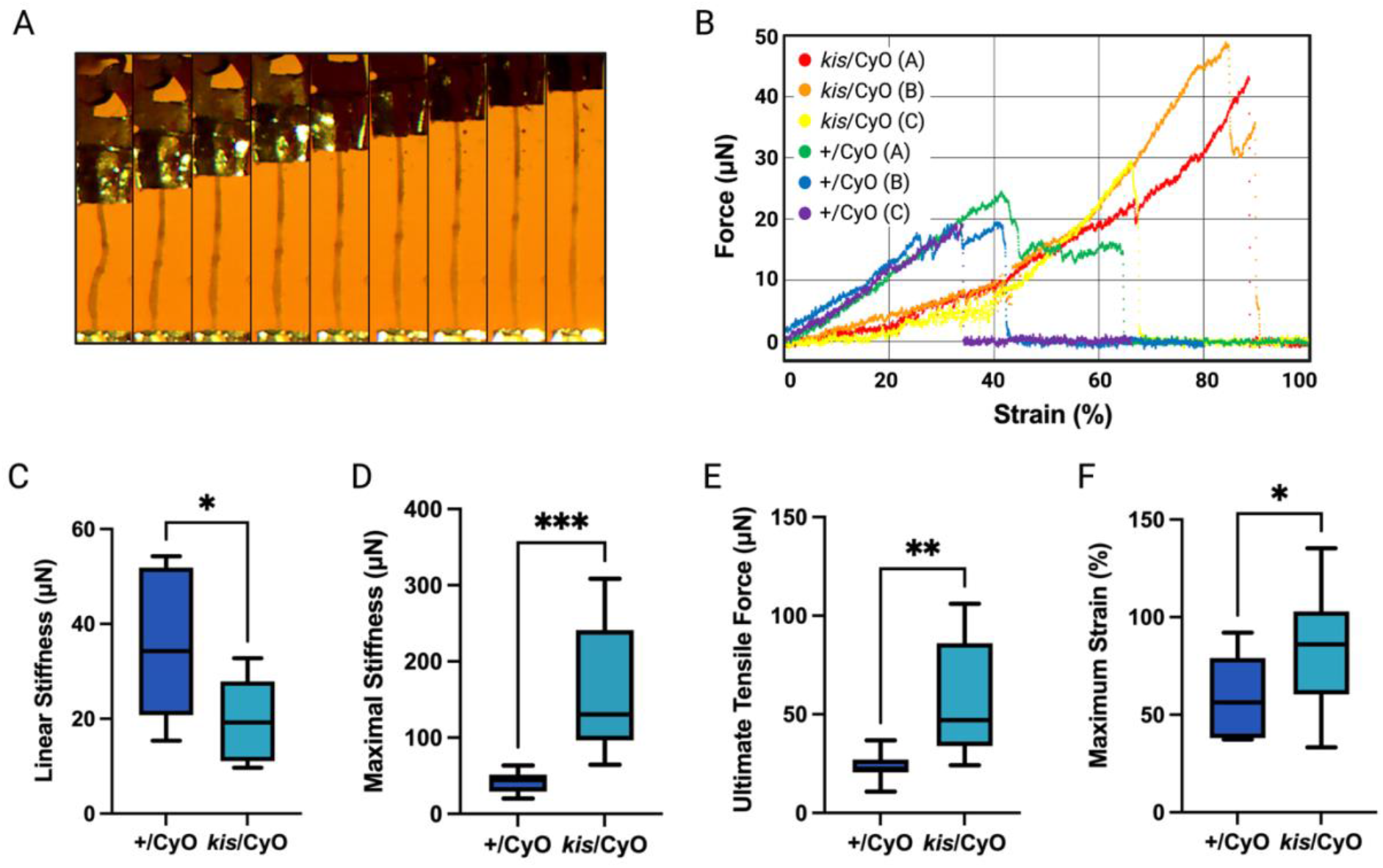
Kismet alters midgut tissue elasticity and tensile strength. (A) Example microscopy image time lapse of a dissected midgut from a *kis*/CyO fly undergoing mechanical testing. Guts were mounted between two metal clips (visible in the top and bottom of the images) and stretched at a constant rate of 0.01 mm/s. Images are 5 mm in height and 20 seconds apart. (B) Six example data sets of dissected midguts from control (+/CyO) flies and mutant (*kis*/CyO) flies. The slope of each curve indicates the stiffness of that sample. The tensile strength is quantified as the highest force and strain the tissue reached before breaking. (C) *kis/CyO* flies had significantly softer midguts when unstretched, compared to control (+/CyO) flies. (D) In contrast to the control, midguts from *kis/CyO* flies strain stiffened substantially, and were significantly stiffer than midguts from +/CyO flies when under strain. (E) Midguts from *kis*/CyO flies also withstood a significantly higher force. (F) Similarly, the *kis*/CyO midguts also reached a higher strain before breaking. In (C) - (E),Student’s *t-*test; * p < 0.05; ** p < 0.01; *** p < 0.001; n = 10 for +/CyO, n = 10 for *kis*/CyO.

### Metagenomic 16S rRNA sequencing

Male flies aged 1-7 days post-eclosion were sterilized in 70% ethanol before guts were dissected in sterile PBS. The foregut was removed by cutting immediately posterior to the proventriculus and the hindgut was removed by cutting immediately anterior to the Malpighian tubules. Midguts (81 from each genotype) were then separated into anterior and posterior regions before being immediately snap frozen in an ethanol dry ice bath. Four samples (+/CyO-anterior, *kis*/CyO-anterior, +/CyO-posterior, *kis*/CyO-posterior) were shipped to GENEWIZ (South Plainfield, NJ), for DNA extraction and sequencing of the V3-V4 16S rRNA gene regions. Resulting sequencing data contained several of GENEWIZ’s proprietary forward and reverse primers, which were removed using Cutadapt (v3.4)(40). The following steps were then performed to process the data within the QIIME2 (v.2021.4) workflow (41): (1) all FastQ files were imported into QIIME2; (2) reads aligning to the *Drosophila melanogaster* genome were removed; (3) reverse reads were trimmed at > 190bp, reads were denoised, dereplicated, paired-end reads merged, and chimeras removed using DADA2 producing an amplicon sequence variant (ASV) table (42); (4) a naive Bayes classifier was trained on the V3V4 region of the 16S rRNA genes in the Genome Taxonomy Database (GTDB v.202) and was used to perform taxonomic classification for each ASV (43); (5) phylogenetic trees were constructed; and (6) table was rarefied before calculation of diversity metrics. The ASV table was converted to a frequency table within the QIIME2 workflow and subsequently, heatmaps were produced to identify genus and species level taxonomic differences between the four samples using the qiime2R (v.0.99.6) package (44).

### Antibiotic depletion

Flies with antibiotic-depleted gut microbiota were created by adding streptomycin (STR) at a concentration of 400μg/mL to the standard cornmeal-yeast-agar medium recipe, as previously described (45). To ensure that gut microbiota were depleted, individual guts of adult male flies were homogenized and spread on De Man, Rogosa, and Sharpe (MRS) plates in serial dilutions. For dissection, flies were anesthetized on ice, the outer surface of the flies were sterilized in 70% ethanol, then rinsed in sterile PBS. Plates were incubated at 25°C for 72 hours prior to counting serial dilutions. Sterile PBS was plated as a negative control.

### Courtship analysis

Post-eclosion males were aged in individual isolation chambers for 5-7 days at 25°C in a humidified incubator on a 12-hour light-dark cycle. Canton S virgin females were housed in vials of up to 10 females and aged for 5-7 days. After the aging period, each male was placed with an untreated Canton S female in a courtship chamber and recorded for 10 minutes. The courtship behaviors—orientation, leg tapping, wing extension, licking, attempted copulation, and successful copulation—were scored. The courtship index (CI) was determined by calculating the percent of time males participated in courtship behaviors for the duration of the assay.

### Statistical analyses

Prism 9 (GraphPad, San Diego, CA) was used to perform all statistical analyses. Normality was tested using the Anderson-Darling test. Parametric data was analyzed using Student’s *t*-test. Non-parametric data was analyzed using the Mann-Whitney U test. Figures were prepared using Prism 9 and BioRender.com.

## Results

### Kismet affects gastrointestinal transit time

We first sought to determine if Kismet could impact GI transit time in *Drosophila*. Individuals with CHARGE syndrome and *CHD8*-associated ASD often have reduced gut motility (27, 29). Similarly, studies using zebrafish have demonstrated that *chd8* knockdown results in slower GI transit, a phenotype attributed to a reduction in the number of enteric neurons (27). Because homozygous null *kismet* mutants are embryonic lethal, we examined *Drosophila* with a null allele of *kismet* (*kis^LM27^*, subsequently referred to as *kis*) balanced over the Curly O (CyO) chromosome, which harbors a wild-type copy of *kismet*. Because our experimental fly strain (*kis*/CyO) included the CyO balancer, we used an isogenic control strain with the same balancer chromosome (+/CyO). To determine if Kismet affected the GI transit time in *Drosophila*, control (+/CyO) and *kismet* mutant (*kis*/CyO) flies were administered food containing bromophenol blue. Flies were observed until the presence of a blue fecal deposit was detected (Figure 1A). We found *kismet* mutant flies had a significantly longer GI transit time: *kis*/CyO flies had an average transit time of 154±92 minutes compared to 84±69 minutes for control flies (p = 0.009; Figure 1B). To determine if the different GI transit times might be attributed to changes in midgut length, midguts from control and *kismet* mutant flies were measured from posterior of the foregut to anterior of the hindgut (Figure 1C). The *kismet* mutant midguts had an average length of 3.90±0.10 mm, which was not significantly different from control midguts, 4.15±0.11 mm (p = 0.110; Figure 1D). Thus, the difference in GI transit time in *kismet* mutants cannot be explained by changes in midgut length. Other possible explanations for slower GI transit include impairments in the enteric nervous system, as observed in *chd8* knockdown zebrafish (27); disruptions in regulatory hormones secreted from enteroendocrine cells (46, 47); changes in contractility of associated visceral muscle tissue (48); and/or structural changes in GI-associated extracellular matrices (ECM), including the peritrophic matrix (49), a protective barrier that lines the lumen of the insect gut.

### Biomechanical properties of the midgut are impacted by Kismet

When dissecting guts for length measurements, we noticed a stark difference in the structural integrity of *kismet* mutant midguts. We therefore conducted high-sensitivity force measurements of the dissected fly gut to determine tissue elasticity and tensile strength. After affixing guts between two clips mounted on a high-precision force transducer and length controller, we extended the midgut along its length at a constant rate (Figure 2A). Midguts were predominantly elastic at the extension rates used here, and the tissue exhibited no relaxation behavior when mechanically tested (data not shown). Although we observed no marked difference in the width of the midgut between samples under light microscopy, we did not have the resolution to accurately quantify the cross-sectional area of the hollow gut. We therefore quantified the elasticity of the midgut as the slope of the force-strain curve (Figure 2B). The *kismet* mutant midgut had a linear stiffness of 19.9±2.9 μN, significantly lower than the 34.6±4.6 μN of control midguts (p = 0.015; Figure 2C). However, the *kismet* mutant midgut strain stiffened substantially, whereas the control midgut exhibited little stiffening when pulled. The *kismet* mutant midguts strain stiffened to a maximal stiffness of 158±26 μN, significantly higher than the control midguts, which only exhibited a maximal stiffness of 41.8±4.4 μN(p < 0.001; Figure 2D). The *kismet* mutant midguts also exhibited an increased tensile strength, reaching an ultimate tensile force of 58.5±9.2 μN and a maximal strain of 83.6±9.2 % before failing. Both measurements were significantly higher than midguts from control flies, which failed at a force of 23.7±2.2 μN (p = 0.002; Figure 2E) and a strain of 58.0±6.6 % (p = 0.037; Figure 2F). These biomechanical measurements showed that, when unstretched, *kismet* mutant midguts are softer than midguts from control flies. However, when stretched, mutant midguts exhibit substantial strain stiffening and become significantly stiffer than controls. Thus, whether the *kismet* mutant midgut is stiffer or softer than the control midgut depends on whether it is stretched. The strain stiffening was dramatic; whereas control midguts largely exhibited the same stiffness regardless of how much they were stretched, mutant midguts doubled in stiffness when stretched. Mutant midguts also withstood a significantly higher force and strain before failing (Figure 2).

### Kismet influences the composition of gut microbiota

Because gut microbiota is both sensitive to and can impact a range of physiological factors in gut tissue (50, 51), we decided to characterize the microbial flora in *kismet* mutant midguts. As with all other experiments, control and *kismet* mutant flies were maintained under identical conditions to ensure that any observed differences could be attributed to the *kismet* null allele. We used 16S rRNA metagenomic sequencing to characterize the microbiota of anterior and posterior midgut regions (Figure 3A). Heatmaps were created to visualize the relative abundance of microbiota within the two genotypes at all taxonomic ranks (Figure 3B-D, Supplementary Figure 1). There were stark differences in the microbial compositions of both anterior and posterior regions of *kismet* mutant midguts at every taxonomic level, the most apparent being a deficit of numerous taxa and the consequent decrease in microbial diversity in *kismet* mutant midguts. However, there were also some taxa found in *kismet* mutant midguts that were either not detected in control midguts or detected in lower abundance. This group included the species *Acetobacter aceti*, which was only detected in *kismet* mutant midguts (Figure 3D); *A. aceti* causes gut dysfunction and shortens lifespan in *Drosophila* (52). Similarly, members of the genus *Providencia* were present in higher abundance in *kismet* mutant midguts (Supplementary Figure 1); *Providencia* are opportunistic pathogens known to interfere with immune activity in *Drosophila* (53).

**Figure 3.**
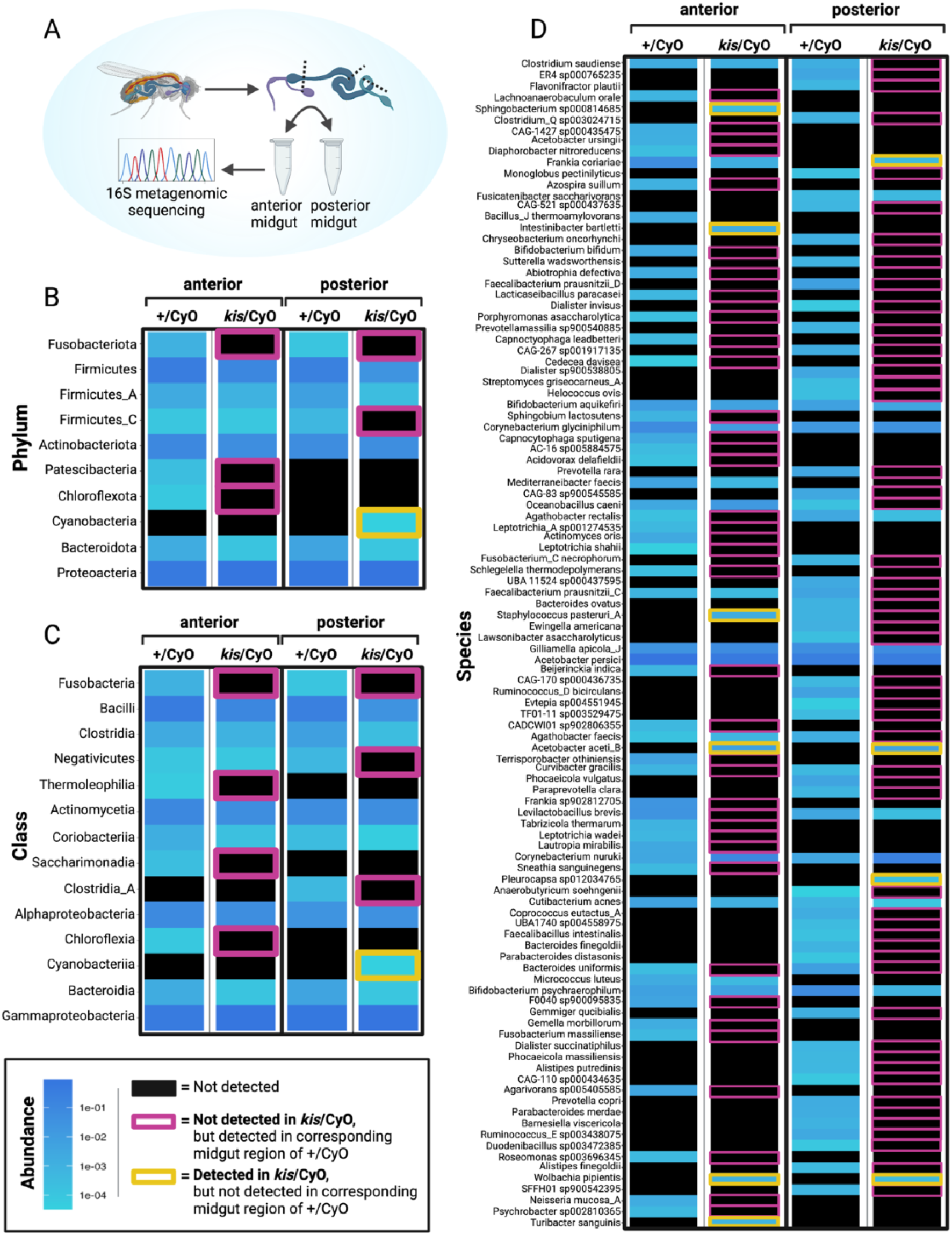
Kismet affects the composition of gut microbiota. (A) Experimental scheme: after removing the foregut and hindgut, midguts were bisected into anterior and posterior midgut regions (dashed lines denote cut sites), which were pooled and used for 16S rRNA metagenomic sequencing. (B - D) Heatmaps indicating the relative abundance of microbial taxa at the (B) phylum, (C) class, and (D) species levels. Relative abundance is reflected in shades of blue. Black indicates taxa that were not detected. Taxa outlined in pink were not detected in the corresponding *kis*/CyO midgut region, but were detected in control (+/CyO) midguts. Taxa outlined in yellow were detected in *kis*/CyO midguts, but were not detected in the corresponding region of control midguts.

Similarly, *Intestinibacter bartletti* was solely detected in *kismet* anterior midguts; while little is known about the role it plays in the fruit fly, a higher abundance of *I. bartletti* has been detected in the guts of children with NDDs (54). Another parallel with NDD-associated gut microbiota was the reduced abundance of butyrate-producing members of the Firmicutes phylum in *kismet* posterior midguts. Multiple studies have found a lower abundance of Firmicutes within the gut microbiome of individuals with ASD (14, 16–18, 55). One of the most abundant butyrate-producing species, *Faecalibacterium prausnitzii,* is also prominently reduced in individuals with NDDs (54); this species was not detected in *kismet* mutant posterior midguts but was present in the control.

### Depletion of gut microbiota differentially impacts courtship behavior

Mutations in *kismet* cause a variety of neurodevelopmental and behavioral phenotypes in fruit flies (31, 34, 35, 37). While the role Kismet plays in neuronal subtypes is understood to affect behavioral phenotypes, we wondered if gut microbiota also affected behavior in *kismet* mutant flies. To address this question, we depleted gut microbiota using food containing low-dose streptomycin (STR) and then compared courtship behavior to flies with unadulterated microbiomes (Figure 4A). To verify that the antibiotic regimen effectively depleted gut microbiota, we measured the colony-forming units (CFU) of STR-treated and untreated flies. By plating homogenized guts, we found that STR significantly reduced the CFUs in midguts from both control (p = 0.001) and *kis*/CyO (p < 0.001) flies (Figure 4B). Next, we examined how depletion of gut microbiota affected courtship behaviors by determining the courtship index (CI), a global courtship score that reflects the fraction of time males spend performing courtship behaviors (56). Depletion of gut microbiota in control flies caused a significant reduction in CI, from a CI of 0.64±0.26 in untreated +/CyO flies to 0.34±0.18 in STR-treated +/CyO flies (p = 0.006; Figure 4C). In contrast, STR-treated *kis*/CyO flies had a significantly higher CI (0.48±0.33) compared to untreated mutants (0.24±0.23; p = 0.021). Therefore, depletion of gut microbiota reduced courtship activity of control flies, but increased courtship activity of *kismet* mutant flies.

**Figure 4.**
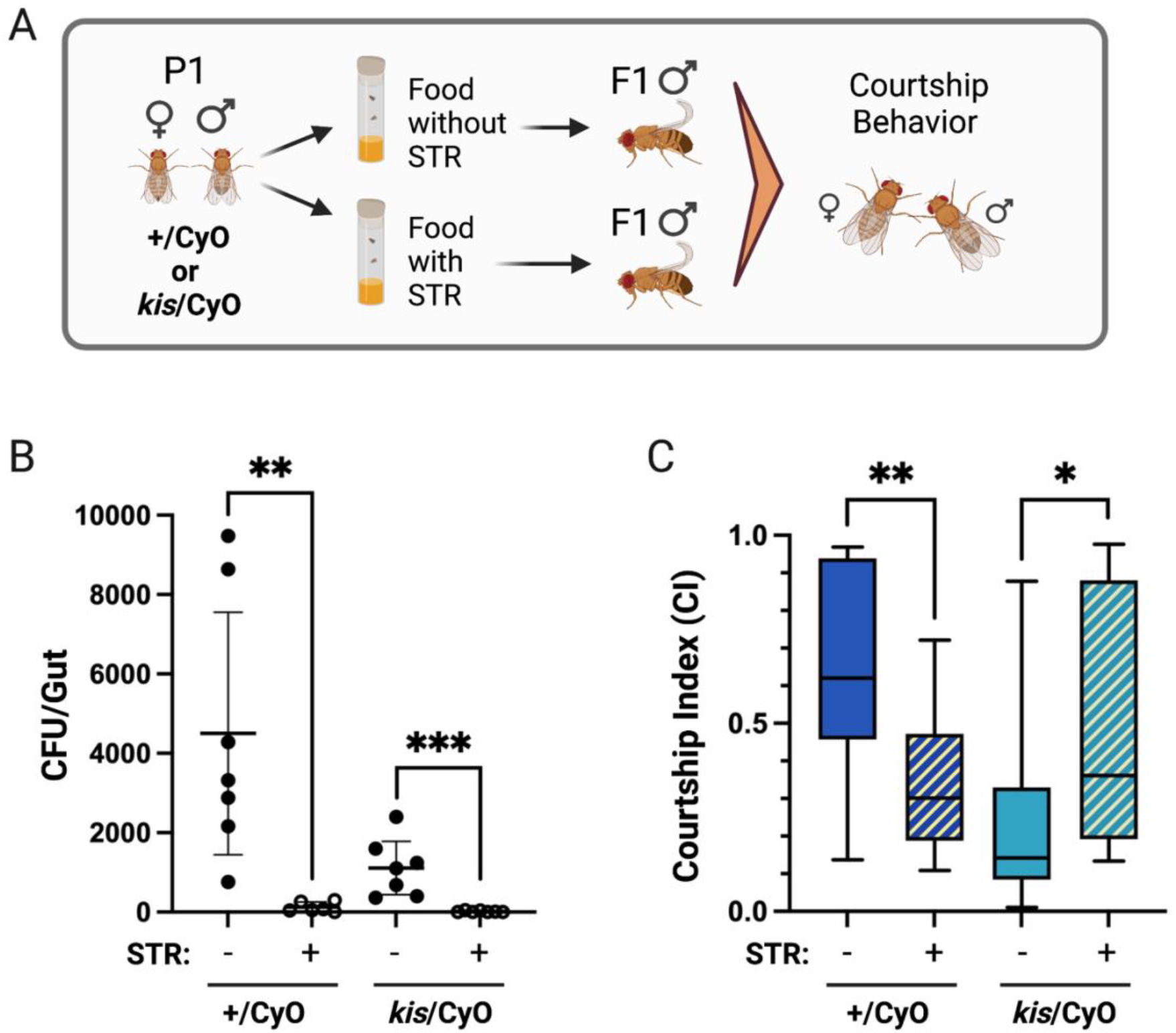
Depletion of gut microbiota differentially impacted courtship behavior in control and *kismet* mutant flies. (A) Experimental scheme: the parental (P1) generation was reared either in control food or food containing streptomycin (STR), as were the first filial (F1) offspring which were used for courtship analyses. Untreated Canton S females were paired with males from each condition. (B) Colony forming units (CFUs) of homogenized whole guts from control (+/CyO) and *kismet* mutant (*kis/CyO*) flies reared in control food (STR -) or STR-containing food (STR +). The horizontal lines indicate means and error bars reflect 95% confidence intervals. Each point represents an individual gut. Mann-Whitney U test; ** p <0.01; *** p < 0.001. (C) Courtship index (CI) of control and *kismet* mutant flies reared in STR- or STR+ food. Mann-Whitney U test; * p < 0.05; *** p < 0.001; n = 12 for each condition.

## Discussion

By investigating gut-related phenotypes of *kismet* mutant *Drosophila,* we identified differences in GI transit rate, biomechanical properties, and microbial composition, as well as a role for the gut-brain axis in modulating *Drosophila* courtship behavior. Given the circuitous nature of the microbiota-gut-brain interactions, we expect there are reciprocal interactions at play that influence each of the observed phenotypes. Because *kismet* encodes a chromatin remodeler that regulates the transcription of many genes within different cell types, the cellular and molecular underpinnings of the observed phenotypes are likely complex and could involve multiple cell types within the brain and gut.

We expected to observe a reduced GI transit rate in *kismet* mutant flies based on studies of *chd8* knockdown zebrafish and reported GI symptoms of humans with *CHD7/CHD8*-associated NDDs (27), where reduced gut motility is attributed to enteric nervous system (ENS) deficits. The slower GI transit rate of *kismet* mutant *Drosophila* may also be affected by reduced numbers and/or deficient innervation of enteric neurons but could also be influenced by structural and mechanical dissimilarities in the gut. For example, changes in contractility of associated visceral muscle tissue (48) or structural changes in GI-associated extracellular matrices (ECM), including the peritrophic matrix, a protective barrier that lines the lumen of the insect gut (49), could affect GI motility. GI activity may also be influenced by disruptions in regulatory hormones secreted from enteroendocrine cells (46, 47), where *kismet* is known to be expressed (33).

Our observation that flies with disrupted GI function also exhibit changes in their gut tissue mechanics is consistent with previous work, which has demonstrated the connection between GI diseases and mechanical changes in the intestine (57), including changes in stiffness (58). While our experiments do not address the underlying molecular changes in the gut tissue that give rise to the observed mechanical phenotype, the high degree of strain stiffening exhibited by *kismet* mutant guts would be consistent with changes in the mechanics or arrangements of cytoskeletal or ECM filaments. For example, stiffness changes of cytoskeletal filaments have been shown to directly result in softer reconstituted networks that undergo more dramatic strain stiffening and can withstand higher forces and strains before failing (59). In addition, the strain stiffening behavior of collagen networks in ECM can be affected by the morphology and crimp of the individual fibers (60). Finally, the peritrophic matrix is composed of aggregated parallel and antiparallel chitin microfibrils associated with chitin-binding proteins; stress-strain curves of different nanostructured chitin composites display variation in the extent of strain-softening and strain-stiffening, depending on the composition of the structure (61). *Chd8/CHD8* affects the expression of genes related to both the cytoskeleton (62) and ECM (63) in mammalian neural progenitor cells, but it is currently unknown how loss of *kismet* affects cytoskeletal and ECM gene expression in *Drosophila* gut epithelia.

We provide evidence that loss of *kismet* affects *Drosophila* gut microbiota by reducing diversity, increasing abundance of pathogenic taxa, and phenocopying characteristics associated with NDD-related gut microbiota. Our data suggest that depletion of gut microbiota can have differential impacts on courtship behavior that may vary according to the level of gut dysbiosis in the native gut microbiome. Given the extensive disruptions in neurodevelopmental processes in *kismet* mutant flies, we were surprised to observe the antibiotic-mediated increase in their courtship activity. One explanation is that depletion of the *kismet* mutant-associated microbiota protects against exacerbation of neuronal phenotypes by factors that would otherwise be secreted by pathogenic microbiota. Conversely, gut microbiota depletion in control flies may induce neuronal phenotypes similar to those typically found in *kismet* mutant flies. For example, mutations in *kismet* are known to impair axogenesis (31, 34). Likewise, mice that undergo embryogenesis in antibiotic treated dams have impaired axon development (64); thus, it would be interesting to explore how gut microbiota depletion impacts neuronal phenotypes, like axon growth and guidance, in *Drosophila*.

There are contradictory results in the field regarding the influence gut microbiota have on *Drosophila* behavior. Changes in gut flora have been attributed to a variety of behavioral changes in fruit flies, including deficits in social behavior (65), sleep (66), and learning and memory (66), though at least two studies have reported insignificant impacts of gut microbiota on *Drosophila* behaviors, including courtship (67, 68). Based on our findings, one explanation for the lack of consensus in the field could be that behavioral consequences of microbiota depletion are dependent on the composition of the *Drosophila* gut microbiome, which is known to vary widely across genotypes and lab environments (69–71).

Although it is unclear how loss of *kismet* promotes changes in gut flora that influence behavior, the corresponding changes in GI transit time and biomechanical properties of the gut are likely involved. Modifications in the peritrophic matrix composition could account for changes in mechanical properties and potentially explain discrepancies in the gut-brain axis—the peritrophic matrix provides a barrier function (72), so changes in its structure could affect permeability to microbes and their metabolites. While further studies are required to elucidate the reciprocal interplay between mechanics, microbiota, and brain, as well as to examine the biophysical mechanisms involved, we suggest that *kismet*-mediated changes in gut structure, mechanics, and function has important roles in the gut-brain axis paradigm.

## Acknowledgements

We would like to extend our gratitude to our staff for invaluable support, especially Douglas Whited, Gordon Zanotti and Caitlin Fox.

## Notes

### Competing Interest Statement

The authors have declared no competing interest.

